# Stick together: Isolation and characterization of exopolysaccharides producing bacteria from degraded permafrost soils

**DOI:** 10.1101/2024.12.28.630587

**Authors:** Muhammad Waqas, Simona Zelenková, Cordula Vogel, Broder Rühmann, Milan Varsadiya, Patrick Liebmann, Haitao Wang, Olga Shibistova, Tim Urich, Georg Guggenberger, Jiří Bárta

## Abstract

Bacterial exopolysaccharides (EPSs) are the high-molecular-weight polymers secreted into their surrounding that play a crucial role in bacterial survival, environmental adaptation, biofilms formation, and interaction with surrounding matrices. EPSs may also contribute to soil structure development by enhancing soil aggregate stability, promoting soil cohesion, and interacting with soil particles through their diverse functional group constituents and conditioning film characteristics. This study aimed to isolate and characterize potential EPSs-producing bacteria from the active layer of two different hydrological landscape of degraded permafrost soils, and from undisturbed intact permafrost soil. A total of 54 bacterial isolates were obtained, representing three phyla: Firmicutes, Actinomycetota, and Pseudomonadota. EPSs production was assessed by determining the polysaccharide content measured as glucose equivalent, and 26 isolates were identified as potential EPSs producers. Among the isolates, *Curtobacterium oceanosedimentum, Frigoribacterium faeni*, *Streptomyces* strains, *Neobacillus bataviensis* and *Mesobacillus subterraneus* exhibited the highest polysaccharide yield. The carbohydrate content of the extracted EPSs varied in both composition and quantity across the different isolates. Uronic acids such as glucuronic acid was found in the EPSs produced by the isolates closely related to *Curtobacterium oceanosedimentum* and *Neobacillus bataviensis*, while the amino hexoses were identified in EPSs extracted from various isolates, including those affiliated with *Bacillus*, *Streptomyces*, *Luteimonas*, and *Phyllobacterium*. The potential EPSs producing bacteria were also found inhabiting in the different horizons of both degraded permafrost soil and undisturbed intact permafrost soil, by determining their relative proportion within the total bacterial community based on 16S rRNA gene sequences similarities.

## INTRODUCTION

Exopolysaccharides (EPSs) are the main part of the complex mixture of organic biopolymers produced and secreted by a wide range of bacteria into their surroundings (Martino 2018). EPSs are categorized into two major types based on the biosynthesis pathway: capsular EPSs, which are tightly associated with the cell surface forming a capsule, and slime EPSs, which are loosely attached to the cell or completely diffuse into the extracellular environment (Li & Yang, 2007; Sheng et al., 2010). EPSs are high-molecular-weight polysaccharides that can be broadly classified into homopolysaccharides and heteropolysaccharides, depending on their monosaccharide composition (Alvarez-Lorenzo et al., 2013; Monsan et al., 2001). Homopolysaccharides consist of a single type of monosaccharide and include examples such as dextran, curdlan, and cellulose. In contrast, heteropolysaccharides are more structurally complex, incorporating diverse monosaccharides such as glucose, galactose, rhamnose, and glucuronic acids (De Vuyst & Degeest, 1999; Oleńska et al., 2021; Wingender et al., 2001). Their structural properties are influenced by specific sugar linkages, branching patterns, and the presence of functional substituents like pyruvates, succinates, and acetates (More et al., 2014; Navarini et al., 1997).

The diverse monosaccharide composition of EPSs, including glucose, galactose, rhamnose, and mannose, contributes to polymer flexibility, enabling the formation of cross-linked networks with the surrounding charge surfaces through covalent bonds, ionic interactions, and hydrogen bonding (Huang et al., 2022). EPSs are rich in distinct functional groups such as hydroxyl, carboxyl, sulfate, phosphate and amino groups, which play a crucial role in cell attachment, and can make complexes with surrounding particles. These functional groups contribute to the anionic or cationic nature of EPSs molecules, enabling electrostatic interactions and hydrogen bonding with various surfaces, including minerals and organic matter (Flemming et al., 1996; Ha et al., 2010). The structurally diverse composition of EPSs along with functional groups, highlights the ecological significance of EPSs- producing bacteria in soil, where they serve a multitude of purposes primarily associated with the biofilm formation and maintenance (Balducci et al., 2023), protecting bacterial cells under various environmental stress, adhesion of the cells to each other, and biofilm attachment to the surface (Wingender et al. 1999; Sandhya and Ali 2015) cryoprotective potentials (Dubey & Jeevaratnam, 2015).

In harsh environments, such as permafrost soils that remain frozen for two or more consecutive years, the unique properties of EPSs are particularly vital, as they facilitate bacterial survival under extreme cold, nutrient scarcity, and fluctuating water availability. Along with it, EPSs are also considered as the main component in the formation and stabilization of soil structures due to their chemical interactions and gelling characteristics(Costa et al., 2018). Therefore, EPSs released by bacteria may have role in protection of soil aggregates under the degradation of permafrost. However, the composition and activity of bacterial communities in permafrost soils are significantly influenced by permafrost degradation (Deng et al., 2015; Jansson & Taş, 2014; Liu et al., 2019; Steven et al., 2008). When rising in Arctic temperatures triggers an increase in the depth of above ground so called active layer, leading to the degradation of permafrost soils and creating conducive environment for bacterial activity (Knoblauch et al. 2013; Koven et al. 2013). This degradation results in several emerging landscape types from which two distinct hydrological conditions, such as, wet soils, e.g. in areas with high moisture, due lower slope position and poor drainage, and dry soils with better drainage and higher evapotranspiration (Jorgenson and Osterkamp 2005; Jorgenson et al. 2013; Liebmann et al. 2024). These landscapes exert different influences on bacterial community compositions and activities, primarily by promoting the decomposition of exposed organic carbon that was previously preserved in frozen ground (Koven et al. 2013). This exposure enhances nutrient availability, which may facilitate bacterial growth, development, and protection through the high production of EPSs under these two different conditions. One way to gain a better understanding of the physiology and ecology of EPSs-producing bacteria is to isolate them from their natural habitats, determine their abundance within the complex soil microbiome, and characterize their EPSs-producing capabilities under laboratory condition. In this study we aimed 1) to isolate and identify the EPSs-producing bacteria from the undisturbed permafrost soil and two degraded permafrost landscapes under lower and higher moisture regimes, respectively, 2) to determine the produced polysaccharides from individual bacterial isolates and 3) to determine the proportion of these EPSs-producing isolates in the total bacterial community.

## MATERIALS AND METHODS

### Soil sample collection

Soil samples were collected from two different landscapes with degraded permafrost and from an undisturbed intact permafrost soil (hereafter, dry, wet and intact sites) located near Fairbanks, Alaska, USA. The detailed descriptions of the studied sites are reported in Liebmann *et al*. (2024). For each site, a soil profile was prepared with a depth of 100 cm for the dry and wet sites, while for the intact site, the profile was excavated until the permafrost table was reached in 45-55 cm soil depth. Soil samples were collected per horizon including organic layer, topsoil and subsoil by taking the material at various depth within each horizon. All samples were constantly kept at -20°C before processing. For the genetic material extraction, approx. three g of soil sample were immersed with two volumes of LifeGuard Soil Preservation Solution (Qiagen, Hilden, Germany) immediately after sampling. All Lifeguard-treated samples were constantly kept at 4°C until processing.

### Isolation of bacteria

For the isolation of bacteria, one g of frozen soil from the topsoil of each site was serially diluted in sterile saline solution (0.9% NaCl) in a total volume of 1 ml. 100 μl of 10^-2^ to 10^-4^ dilutions were plated on three distinct culture media. Standard I (1 g glucose L^-1^, 15 g peptone L^-1^, 3 g yeast extract L^-1^, and 6 g NaCl L^-1^), Reasoneŕs 2A agar (Reasoner & Geldreich, 1985) and Nutrient agar (2 g Yeast extract, 5 g Peptone L^-1^, 1g Lab-Lemco L^-1^ and 5 g NaCl L^-1^) were utilized to ensure the recovery of a diverse range of bacteria, including general heterotrophic, slow-growing oligotrophic bacteria, and actinomycetes. The plates were incubated at 22°C for 48–72 hours. The 22°C was chosen based on higher CFU (data not shown). After the incubation, well-grown single colonies were picked from each diluted agar plate and streaked onto corresponding fresh agar plates, followed by incubation. The streaking process was repeated a minimum of three times to ensure the isolation of pure colonies. The pure colonies were subsequently grown in the Nutrient liquid medium for DNA isolation.

### Screening soil isolates for EPSs production

For screening of EPSs production, soil isolates were streaked on agar medium (EPS medium) specifically designed exopolysaccharides production described in Rühmann et al. (2015). The plates were incubated at 22°C for 48–72 hours. The isolates that were able to grow on EPS medium and formed a mucoidal/slimy colony were considered as potential EPSs-producers. They were further screened for exopolysaccharides production via automated analytical system involves various polysaccharide detection modules, including determination of total glucose content using phenol-sulfuric-acid transformation, and carbohydrate content by High-throughput-1-phenyl-3-methyl-5-pyrazolone (HT-PMP)-derivatization of the carbohydrates coupled to Electrospray Ionization Mass Spectrometry. as described in Rühmann et al. (2016). The production of polysaccharides measured as glucose equivalent by soil isolates exceeding 100 mg L^-1^ were considered as the greater EPSs-producers. This threshold was set to distinguish between isolates with high and low EPSs production, making it easier to focus on those with significant activity.

### DNA extraction and Polymerase Chain Reaction

The extracted DNA from soil samples and isolates were quantified by Quantus fluorometer (Promega BioSystems, Sunnyvale, California, USA). For this, 2 μl of DNA sample was added to to 98 μl 1x TE buffer and 100 μl QuntiFluor dye (QuntiFluor dsDNA dye, Promega Corporation, Madison, USA) in a 1 ml clean Eppendorf tube. A blank and standard were also prepared by mixing 100 ul of 1x TE buffer with 100 μl of QuantiFluor dye or standard DNA provided by manufacturer, respectively. The tubes were incubated in the dark for 10 minutes and the fluorimeter was calibrated by standard and blank samples before quantifying the DNA samples.

Genomic DNA extracted from isolates was utilized to amplify the 16S rRNA gene using master mix composed of DNA, ddH_2_O, PCR buffer, BSA and primers: 9bfm (forward primer: 5’- GAGTTTGATYHTGGCTCAG-3’) and 1512uR (reverse primer: 5’-ACGHTACCTTGTTACGACTT-3’). For a positive control DNA of *Escherichia coli* (strain ATCC 9637) and for a negative control H_2_O were used. The PCR amplification was performed using the PCR thermal cycler (SensoQuest GmbH, Germany) with the following protocol: initial denaturation for three minutes at 95°C, followed by 30 cycles of each denaturation for one minute at 95°C, annealing for one minute at 52°C and elongation for 90 seconds at 72°C. The final elongation was for 10 minutes at 72°C. After the amplification, 4 μl of PCR product was determined by electrophoresis on 1% agarose gel and visualized by trans illuminator (Azure 200, Azure Biosystem, Inc, US). The length of the band was compared to 1 kb DNA ladder and the correct PCR products were then sent to SEQme sequencing company for Sanger sequencing. The obtained sequence data underwent quality trimming and the forward and reverse reads were joined using Geneious Prime bioinformatics software (Java Version 11.0.18+10 (64 bit). The nucleotide sequences were then searched in 16S rRNA gene NCBI database by standard nucleotide BLAST.

### Determination of proportion of EPSs isolates in total community

The proportion of the EPSs-producing soil isolates in the total relative proportion of bacterial community from different horizons of each landscape was determined. For this, 16S rRNA gene sequences from the EPSs-producing soil isolates were used as queries to search against a local 16S rRNA gene database constructed from the representative sequences of zero-radius operational taxonomic unit (zOTUs) from the total bacterial community using BLAST (Figure S1, Supporting Information). Afterward, a subset of assigned zOTUs sharing sequences with more than 97% similarity with 16S rRNA sequence of pure isolates was used to calculate their proportion within the total bacterial community across the different horizons within all the three studied sites.

The relative proportion of the bacterial community was calculated by dividing the count of each zOTUs by the sum of counts within each sample and further summarized across the sites and horizons. The data were filtered to include only the phylum level and the mean relative proportions were then determined. The proportion of EPSs-producing soil isolates among the total bacterial community was calculated by taking the subset of assigned zOTUs with their relative proportions. The data were then filtered at genus level and computed to mean relative proportion. All the figures were generated using the ggplot v 3.4.4 package (H, 2016).

## RESULTS

### Isolation and characterization of potential EPSs-producing bacteria

The bacterial isolation was carried out using three distinct culture media, resulting in the isolation of 17, 21, and 16 bacterial isolates from soil of the dry, wet and intact sites, respectively (Table 1, TableS1, Supporting Information). These isolates were grown on specific polysaccharide production medium for the formation of mucoidal “sticky” colonies, which served as an indicator of their potential to produce EPSs. The majority of soil isolates exhibited growth in the form of mucoidal colonies on medium and signifying their potential as EPSs producers (Table S1, Supporting Information). Out of 54 isolates, 48 were successfully sequenced and classified into three phyla: Firmicutes, Actinomycetota, and Pseudomonadota. Within the Firmicutes, potential EPS producing isolates obtained from all three sites were taxonomically assigned to several genera, including *Bacillus, Peribacillus, Neobacillus,* and *Mesobacillus*. Six isolates were identified as *Bacillus mycoides*, while other six isolates from the wet site were assigned to *Bacillus subtilis*, *Peribacillus simplex*, *Neobacillus bataviensis* and *Mesobacillus subterraneus* (Table 1). Among the Actinomycetota, five isolates from the intact site were assigned to *Frigoribacterium faeni*, *Curtobacterium oceanosedimentum* and *Pseudarthrobacter sp.* and three isolates from the dry site were related to *Microbacterium flavescens* and *Streptomyces sp*. One isolate from the wet site was assigned to *Streptomyces luozhongensis*. Within the Pseudomonadota, isolates were taxonomically classified to *Phyllobacterium sp.* and *Luteimonas arsenica* (Table 1). We found only 5.6% overlapping with one similar isolate such as *Bacillus mycoides* between soil of the dry, wet, and intact sites and another one common isolate such as *Curtobacterium oceanosedimentum* found in soil of both wet and intact sites (Fig. S2, Supporting Information).

**Table 1.** Taxonomic classification of soil isolates identified as potential EPSs producers, that produced more than 100 mg L^-1^. Isolates sharing more than 97% sequence similarity are presented at the species level and lower than 97% are presented at family level. Polysaccharide contents are listed for isolates exceeding 100 mg L^-1^ production

| Phylum | ID | Closest Blast Hit 16s rRNA sequencing | % Identity | Locality | Polysaccharide content (mg L<sup>-1</sup>) |
| --- | --- | --- | --- | --- | --- |
| **Firmicutes** |  |  |  |  |  |
|  | S29 | *Bacillus mycoides* strain ATCC 6462 | 99.9 | Intact | 145.34 |
|  | S31 | *Bacillus mycoides* strain ATCC 6462 | 100 | Intact | 180.69 |
|  | S37 | *Bacillus mycoides* strain ATCC 6462 | 97.6 | Intact | 150.05 |
|  | A6 | *Bacillus proteolyticus* strain MCCC 1A00365 | 98.0 | Intact | 170.17 |
|  | A13 | *Bacillus mycoides* strain ATCC 6462 | 100 | Dry | 212.37 |
|  | S28 | *Bacillus mycoides* strain ATCC 6462 | 100 | Dry | 140.79 |
|  | A24 | *Bacillus mycoides* strain ATCC 6462 | 100 | Wet | 515.24 |
|  | S38 | *Bacillus subtilis* strain SBMP4 | 99.2 | Wet | 171.48 |
|  | S39 | *Bacillus subtilis* strain SBMP4 | 98.0 | Wet | 164.54 |
|  | S3 | *Peribacillus simplex* strain LMG 11160 | 96.5 | Wet | 100.0 |
|  | A25 | *Neobacillus bataviensis* strain NBRC 102449 | 99.0 | Wet | 1148.6 |
|  | S19 | *Neobacillus pocheonensis* strain Gsoil 420 | 99.1 | Wet | 146.0 |
|  | S6 | *Mesobacillus subterraneus* strain COOI3B | 98.2 | Wet | 911.44 |
| **Actinomycetota** |  |  |  |  |  |
|  | S17 | *Frigoribacterium faeni* strain 801 | 99.4 | Intact | 1309.14 |
|  | S18 | *Frigoribacterium faeni* strain 801 | 99.5 | Intact | 735.79 |
|  | A17 | *Curtobacterium oceanosedimentum* strain ATCC 31317 | 99.3 | Intact | 1309.14 |
|  | S15 | *Pseudarthrobacter sulfonivorans* strain ALL | 98.6 | Intact | 115.63 |
|  | S23 | Micrococcaceae | 91.7 | Intact | 208.21 |
|  | A11 | *Streptomyces brevispora* strain BK160 | 99.5 | Dry | 342.92 |
|  | A2 | *Microbacterium flavescens* strain 401 | 98.3 | Dry | 136.85 |
|  | S9 | *Streptomyces luozhongensis* strain TRM49605 | 99.5 | Wet | 118.63 |
|  | S7 | *Curtobacterium oceanosedimentum* strain ATCC 31317 | 98.8 | Wet | 216.69 |
| **Pseudomonadota** |  |  |  |  |  |
|  | A12 | *Luteimonas arsenica* strain 26-35 | 99.8 | Dry | 227.26 |
|  | A22 | *Phyllobacterium loti* strain S658 | 99.0 | Wet | 177.28 |
|  | A23 | *Phyllobacterium endophyticum* strain PEPV15 | 99.5 | Wet | 124.15 |
|  | S5 | Enterobacteriaceae | 91.3 | Wet | 667.23 |

### Differences in EPSs production by soil isolates

In general, isolates with production of polysaccharide measured as glucose equivalent exceeding 100 mg L^-1^ were considered as potential EPSs-producing isolates (Table 1 and Fig. 1). Two isolates showing similarity to *Curtobacterium oceanosedimentum* and *Neobacillus bataviensis* yielded the highest polysaccharide more than 1000 mg L^-1^ such as 1309 mg L^-1^ and 1149 mg L^-1^ respectively. This was followed by the *Frigoribacterium faeni* and *Mesobacillus subterraneus* exhibited polysaccharide above 900 mg L^-1^. Other isolates belonged to Firmicutes, such as *Bacillus subtilis*, *Peribacillus simplex*, and *Bacillus mycoides* yielded polysaccharide upto 200 mg L^-1^, except for the one isolate related to *Bacillus mycoides* (A24) accounted 515 mg L^-1^ polysaccharide content. Similarly, isolates assigned to *Streptomyces sp.*, *Pseudarthrobacter sulfonivorans* and *Microbacterium flavescens* yielded in range of 100 to 350 mg L^-1^. From the phylum Pseudomonadota, isolate of family Enterobacteriaceae exhibited the polysaccharide upto 667 mg L^-1^, *Luteimonas arsenica* 227 mg L^-1^ and *Phyllobacterium loti* 177 mg L^-1^. In total, five isolates from soil of the dry site, 12 isolates from soil of the wet site, and nine isolates from soil of the intact site were identified as potential high EPSs-producers, as their polysaccharide equivalent as glucose surpassed 100 mg L^-1^.

**Figure 1.**
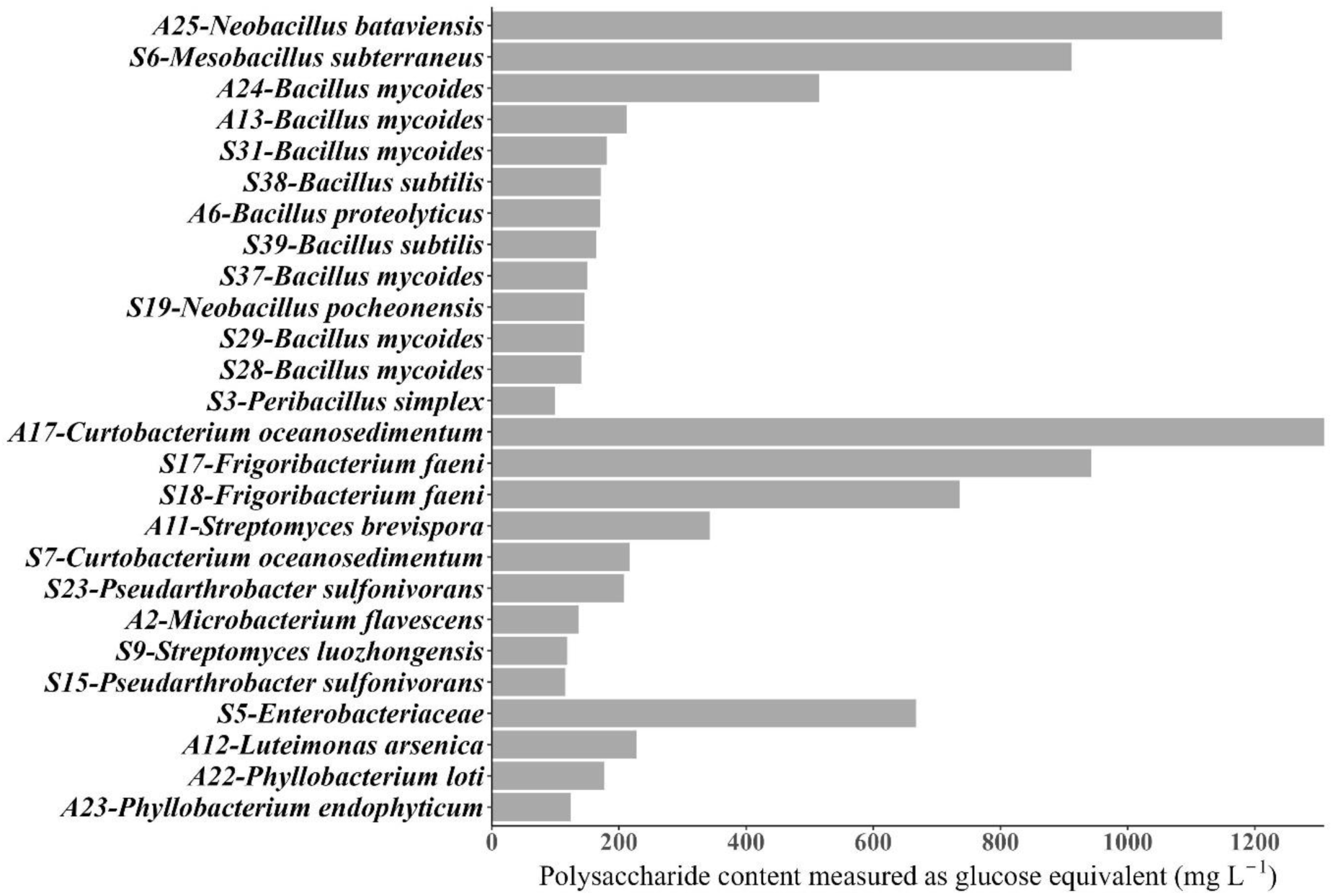
A: Polysaccharide production measured as glucose equivalent in mg L^-1^.

### Carbohydrate contents of EPSs produced by soil isolates

The soil isolates identified through 16S rRNA sequencing revealed a diverse array of EPSs enriched with various carbohydrate derivatives determined by High-throughput-1-phenyl-3-methyl-5-pyrazolone (HT-PMP)-derivatization of the carbohydrates coupled to Electrospray Ionization Mass Spectrometry (Table 2). Among the Firmicutes isolates, *Bacillus mycoides* exhibited a substantial presence of hexoses, particularly glucose (Glc) (up to 65 mg L^-1^), mannose (Man) (upto 5 mg L^-1^) and galactose (Gal) (upto 102 mg L^-1^), along with different levels of amino hexoses, such as glucosamine (GlcN) (up to 62 mg L^-1^), Galactosamine (GalN) (upto 10 mg L^-1^) and N-acetylglucosamine (GlcNAc) (upto 12 mg L^-1^). One isolate of *Bacillus mycoide* (A24) with exception of carbohydrate derivatives in their EPSs were composed of only Gal (102 mg L^-1^) and Man (3 mg L^-1^). The sum of the sugar content in EPSs produced by isolates of *Bacillus mycoide* was varying 52 to 155 mg L^-1^ (Fig. 2). The isolate *Neobacillus bataviensis* and *Mesobacillus subterraneus*, that exhibited the highest EPSs among the Firmicutes showed distinct carbohydrate profiles. For examples, the EPSs produced by *Neobacillus bataviensis* was composed of Glc (103 mg L^-1^) and glucuronic acid (GlcUA) (21 mg L^-1^), along with smaller amounts of Gal (2 mg L^-1^), rhamnose (Rha) (6 mg L^-1^), and GlcN (4 mg L^-1^). The EPSs produced by *Mesobacillus subterraneus* was characterized by high Glc content (203 mg L^-1^) and Man (29%), Rha (12 mg L^-1^), and GlcN (6 mg L^-1^). The sum of the carbohydrate content was higher for the *Mesobacillus subterraneus* (249 mg L^-1^) than the *Neobacillus bataviensis* (135 mg L^-1^). The carbohydrate contents of the EPSs produced by isolates belonged to Actinomycetota were high in the amount of Glc followed by the Man and GlcN. The highest EPSs producers such as *Curtobacterium oceanosedimentum* showed a carbohydrate profile dominated by Glc (114 mg L^-1^) and glucuronic acid (GlcUA) (33 mg L^-1^), with minor amounts of Gal (1 mg L^-1^) and Rha (7 mg L^-1^). GlcUA was only found in the EPSs of *Curtobacterium oceanosedimentum,* and was not detected in the EPSs of other isolates of Actinomycetota. *Microbacterium flavescens* on the other hand, Man (12 mg L^-1^) and Rha (7 mg L^-1^) were only detected in their produced EPSs. Similarly, the sum of the carbohydrate derivatives was higher for the isolates of *Frigoribacterium sp.*, followed by the *Curtobacterium oceanosedimentum*, and the least was detected in the EPSs of *Microbacterium flavescens*. In the Pseudomonadota group, EPSs from *Luteimonas arsenica* was of only hexoses, in which the Gal was the highest in amount (125 mg L^-1^). While the EPSs of other isolates of Pseudomonadota were more enriched with Glc, GlcN and GlcNAc. The sum of carbohydrate content was about 120 to 190 mg L^-1^ for the isolates of Pseudomonadota.

**Figure 2:**
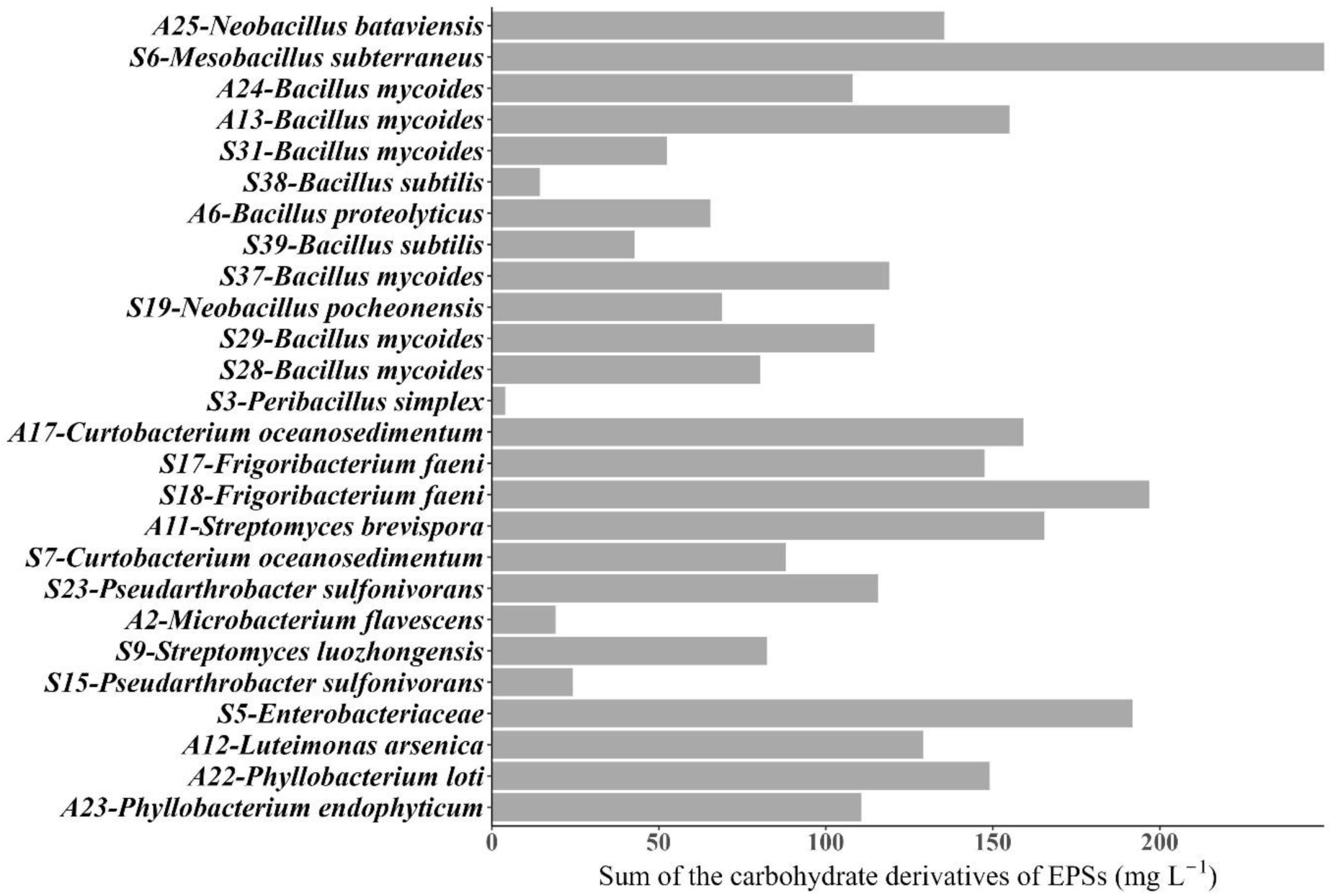
Sum of the sugar derivatives determined from the extracted EPSs.

**Table 2:**
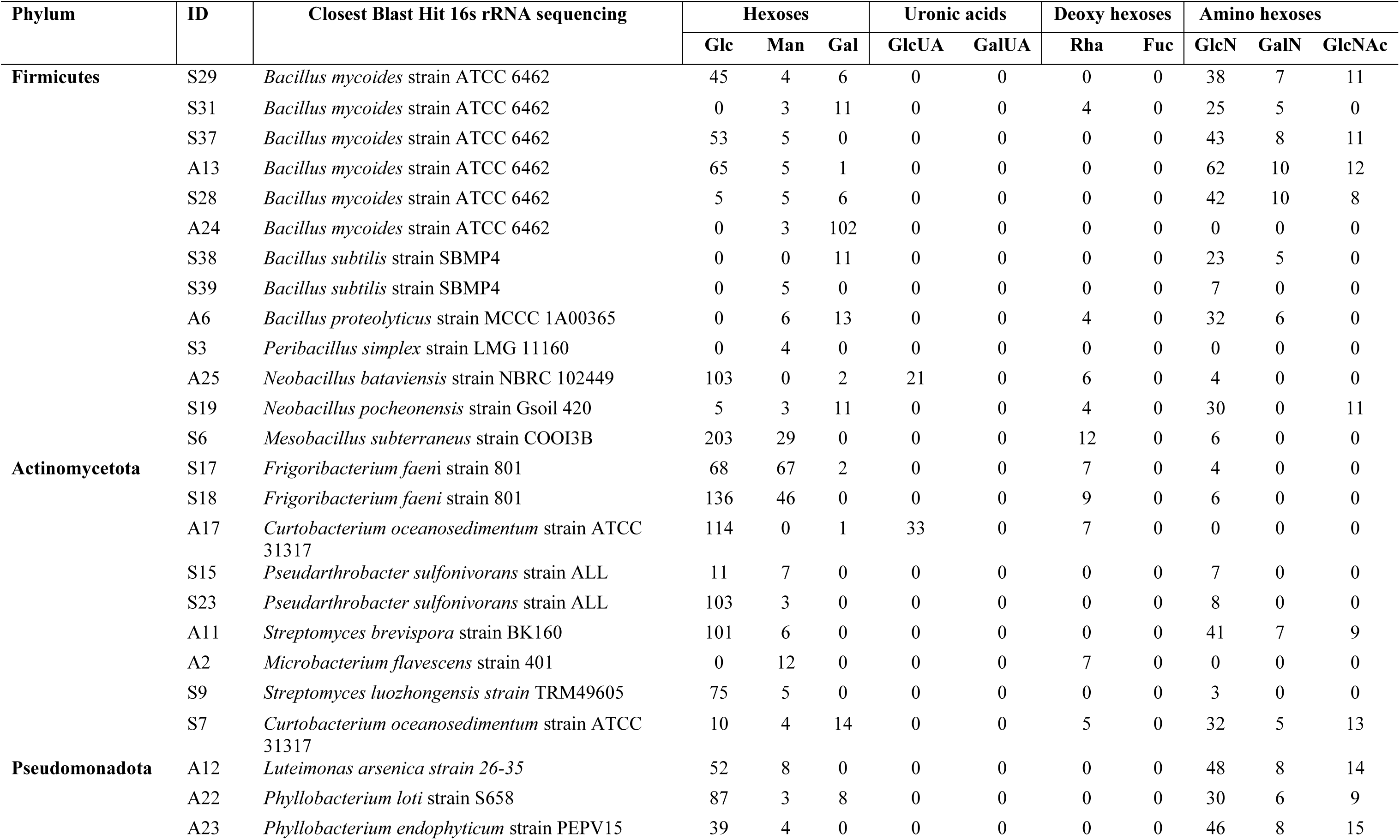
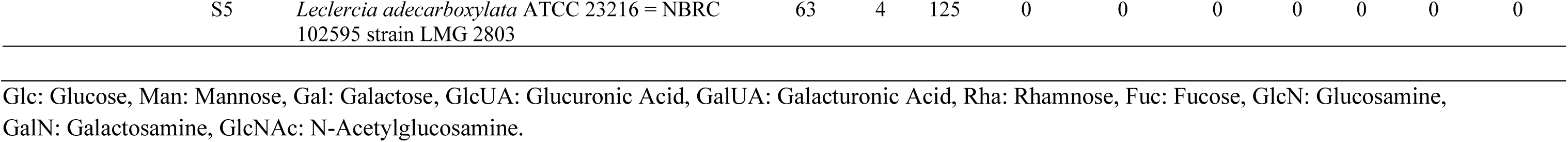
Sugar derivatives of EPSs produced by soil isolates measured in mg L^-1^

### Proportion of EPSs-producing isolates in total bacterial community

The 16S rRNA gene sequence of 24 EPSs-producing isolates showed similarities higher than 97% to the sequence of 13 zOTUs (Table S2). The proportion of assigned zOTUs within the total bacterial community was different among the three sites as well as within the horizon (Fig. 3). The potential EPSs-producing isolates represented approx. 2% of total bacterial community in soil of the dry site, ∼0.5% in soil of the wet site, but below 0.36% in the soil of the intact site. Among the Firmicutes, *Bacillus subtilis* (zOTU3) was predominantly detected in highest proportion in both topsoil (∼0.67%) and subsoil (∼0.39%) of the dry site. However, it was almost undetected in the horizons of both the other sites (intact and wet sites), with levels below 0.01%. In contrast, *Bacillus mycoides* (zOTU2010) was only found with proportion of 0.03% in the topsoil of the wet site. *Peribacillus simplex* (zOTU2176) was the second after *Bacillus subtilis* found in high proportion of 0.24% in the topsoil of dry site, while it was not detectable in the subsoil. Additionally, their proportion in the topsoil of the wet site was 0.03% and was also detected to have less proportion in the organic layer and subsoil of both intact and wet site. Similarly, *Neobacillus bataviensis* and *Neobacillus pocheonensis* assigned to zOTU702 and *Mesobacillus subterraneus* to zOTU3623 were observed in proportions of 0.14% and 0.02% respectively, in the topsoil of the wet site, contrasting with their proportions of less than 0.001% in other horizons across all three sites (Fig. 3).From the Actinomycetota, two assigned zOUTUs, *Streptomyces luozhongensis* (zOTU1852) and *Streptomyces brevispora* (zOTU1848) were found in high proportions in the organic layer of the dry site of 0.36% and 0.1%, respectively, while the proportion of *Curtobacterium oceanosedimentum* (zOTU2238) was high in the organic layer of the intact site with 0.14%. Moreover, *Frigoribacterium faeni* (zOTU1136), *Pseudarthrobacter sp.* (zOTU1072) and *Microbacterium flavescens* (zOTU1055) shared almost similar proportion across the topsoil and subsoil in all three sites, with the exception of *Pseudarthrobacter sp.*, which was also found in the organic layer of the dry site (Fig. 3).The *Phyllobacterium endophyticum* of Pseudomonadota assigned to zOTU3875 exhibited a range of high proportion of 0.03% in the topsoil to a low proportion of 0.001% in subsoil of the wet site, while it was not fund in any horizons of the dry and intact sites. Lastly, *Luteimonas arsenica* (zOTU2611) was also found in all horizons except organic layer of dry site and subsoil of intact site.

**Figure 3.**
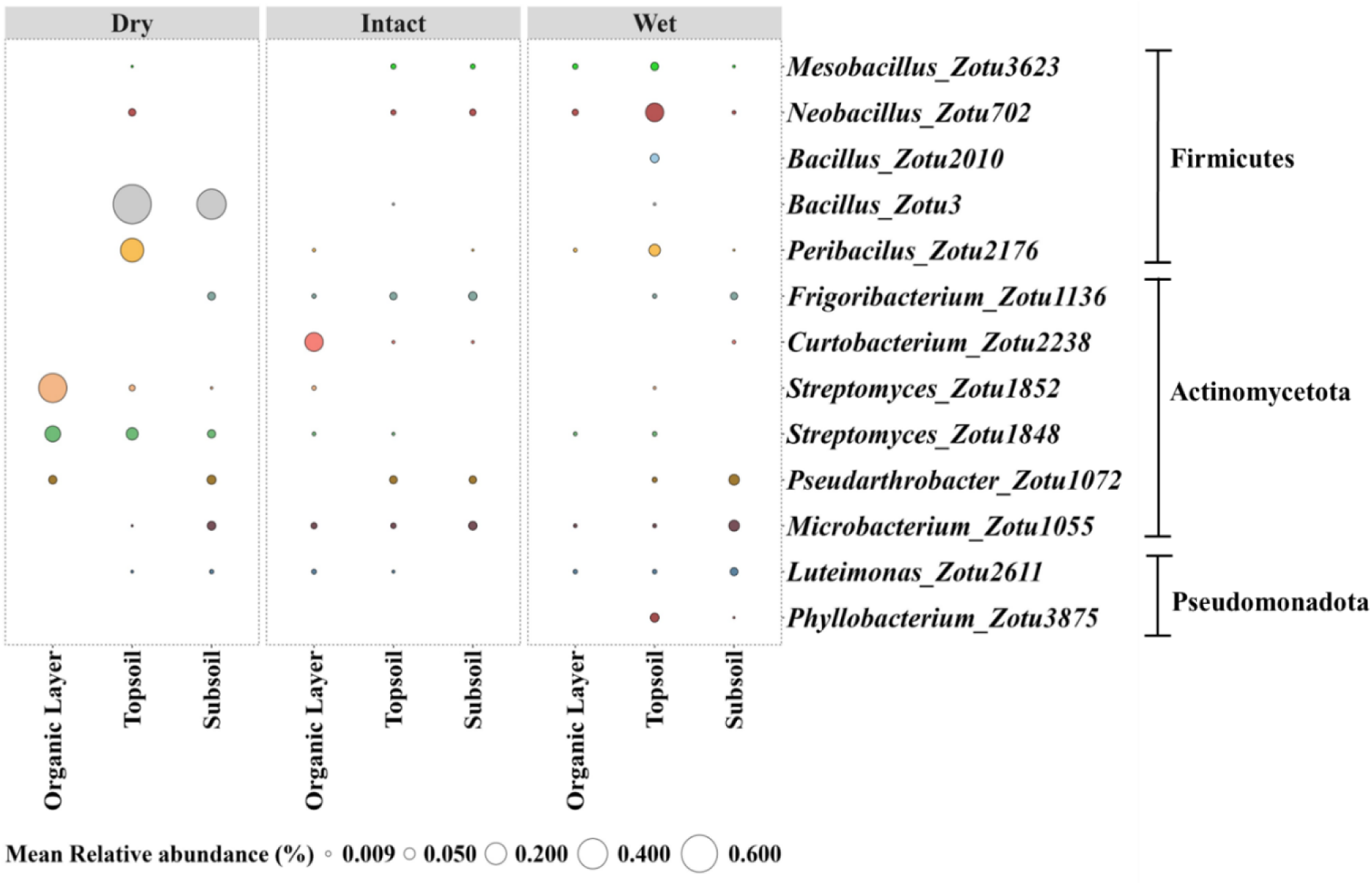
Relative proportion of assigned zOTUs for EPSs-producing soil isolates at genera level in total bacterial community of two sites with degraded permafrost soil (dry and wet site) and site with undisturbed permafrost soil (Intact site).

## DISCUSSION

### Bacterial exopolysaccharides and their carbohydrate content

Bacterial EPSs have long been recognized for their structural diversity, which provides a wide range of physicochemical and biological properties. The structural properties of EPSs serve various roles, including concealing the bacterial surface, acting as adhesives for interactions with other bacterial surfaces or substrates, stabilizing biofilm structures, functioning as signaling molecules and providing protection against environmental stresses (Flemming & Wingender, 2010; Wingender et al., 1999). We utilized the cultural base approach together with further screening for exopolysaccharide production and carbohydrate contents using an automated analytical system. Based on it, 24 bacterial strains were selected as potential EPSs producers out of 54 isolates obtained from the dry and wet permafrost degradation landscapes and undisturbed intact permafrost soil. Two isolates, related to *Curtobacterium oceanosedimentum* and *Neobacillus bataviensis*, were the only strains to exhibit high EPSs production containing uronic acids, distinguishing them from the other isolates. EPSs typically contain neutral carbohydrates, such as hexose and pentose, along with uronic acids like glucuronic and galacturonic acids (More et al., 2014). The functional anionic groups in these uronic acids provide a high cation exchange capacity and enable strong complexation with metal ions (Guibaud et al., 2003; Ha et al., 2010). This binding of EPSs to divalent cations like Ca²⁺ and Mg²⁺ promotes the structural integrity of bacterial aggregates, a key interaction in biofilm formation and stability (Mayer et al., 1999). Additionally, amino hexoses component of EPSs such as glucosamine and galactosamine contain amino groups that interact with surrounding molecules, such as metals and organic compounds (Lahiri et al., 2022; B.-B. Wang et al., 2018). These amino hexoses were identified in EPSs extracted from various isolates, including those affiliated with *Bacillus*, *Streptomyces*, *Luteimonas*, and *Phyllobacterium* (Table 2).

In environments like permafrost soils, where nutrient availability is limited and bacterial survival is challenged by extreme cold and low water activity, the uronic acids and amino hexoses content in EPSs may play important role in microbial adaptation by stabilizing with surrounding charged particles. Soil particle composition, including clay, silt, and organic matter, may directly influence the interaction of EPSs amino sugars, and uronic acids. Mineral components like aluminum and iron oxides, along with organic matter, provide surfaces for metal ion binding through the formation of form charged sites on their surfaces (Kleber et al., 2015; Wagai et al., 2020), which then complexes with amino sugars like glucosamine and galactosamine, and the uronic acids in EPSs. These interactions enhance bacterial stability, biofilm formation, and nutrient retention within the soil matrix (More et al., 2014), and may also enhances the soil’s overall porosity and aggregate stability (Zhang et al., 2024) and conserving the soil organic carbon pool from degradation and reducing soil dispersion (Kong et al., 2011). However, the production of EPSs with varying compositions is influenced by external environmental factors such as nutrient availability, moisture, pH and temperature (More et al., 2014), in contrast to the production observed with pure strains under controlled laboratory conditions.

### Proportion of potential EPSs-producing isolate in total community

We employed a bioinformatics approach to assess the proportion of potential EPSs-producing soil isolates within the proportion of total bacterial community present in the different horizons of intact and degraded permafrost soil. The utilization of high-throughput sequencing and bioinformatic tools allowed us to discern the taxonomic affiliations and relative abundance of potential EPSs-producing isolates within the overall microbial community structure (Fig. 3 and Table S2, Supporting Information). The proportion of potential EPSs-producing isolates within total community varied among the three sites, with the community structure exhibited greater diversity in soil of the wet and intact sites compared to the one in the dry site (Fig. 3). However, some of the assigned genera such as *Streptomyces* (zOTU1852 and zOTU1848), *Bacillus* (zOTU3) and *Peribacillus* (zOTU2176) to EPSs producing soil isolates found only or with higher proportion in the different horizon of the dry site (Fig 3). This may be due to the hydrophilic nature of carbohydrates in EPSs due to their hydroxyl groups plays a key role in maintaining bacterial hydration and preventing desiccation and supporting the transport of nutrients in the cold and water-limited conditions of permafrost soils, or even during the freezing period (Ali et al., 2020). Strains of *Streptomyces, Bacillus* and *Peribacillus* are known for their ability to produce EPSs as potential response to low soil moisture level (Wang et al. 2003; Marvasi et al. 2010; Homero et al. 2021) and were also investigated for their role in soil aggregation (Aspiras et al., 1971; Raliya et al., 2014). Furthermore, On the other hand, the assigned genera *Mesobacillus*, *Neobacillus*, *Microbacterium and Pseudarthrobacter* were higher in topsoil and subsoil of the intact and wet sites compared to the dry site, whereas some of them were only found more prominent in the subsoil. Similarly, *Curtobacterium oceanosedimentum* (zOTU2238) was found in wet subsoil and intact throughout all the horizon. The presence of these genera in the wet and intact sites suggests their acclimation to high moisture content and low temperatures through the EPSs production in their surroundings. In permafrost soils, where nutrient is often limited, the hydrophobic regions in certain carbohydrates of EPSs may play a critical role in bacterial survival. These hydrophobic regions help EPSs adsorb organic compounds, including aromatics and aliphatics, that are released from decomposed organic matter (Huang et al., 2022; Ibrahim et al., 2022). By enhancing the binding potential of EPSs, these hydrophobic interactions may assist bacteria in retaining nutrients in environments where organic substrates are scarce or undergoing slow decomposition due to the low temperatures and high temperature.

EPS production, with its varying components can enables bacteria to survive and thrive under diverse environmental conditions. In addition to their role in growth and nutrient retention, EPSs may also contribute to soil structure stability and aggregate formation through the interaction of functional groups and the gelling properties of the EPSs, which could play a key role in maintaining soil cohesion and enhancing aggregate development (Kaci et al. 2005; Vardharajula and Sk 2014; Deka et al. 2019).

## CONCLUSION

Bacteria have adopted unique mechanisms to cope with and survive in cold habitats experiencing environmental fluctuations. One of these adaptations includes the production of cryoprotective agents in the form of EPSs (Krembs et al., 2002). These agents also act as a gluing substance, physically promoting soil aggregation (Costa et al. 2018). The current study provides information about the bacterial strains isolated from two degrading permafrost soils and an undisturbed intact permafrost soil. These isolates are particularly important for EPSs production in order to survive in extreme conditions characterized by low temperatures, varying moisture levels and limited nutrients. We observed a notable tendency for high polysaccharide production with different carbohydrates derivatives by the soil isolates, indicating that these are potentially high EPSs producers. They were also observed in varying proportion within the total bacterial community in both degraded permafrost soils and undisturbed intact permafrost soil. The results of this study suggest that the presence of bacteria in such environment may be due to their production of EPSs, that provide protection from environmental stress. This EPSs production in their nature environment may also influence the soil aggregate formation in different degradation landscapes of permafrost soils.

## DATA AVAILABILITY

Raw fastq 16S rRNA gene sequences of the whole microbiome were uploaded to European nucleotide archive (ENA) under the study n. PRJEB79237. Sequences of the pure strains were uploaded also to ENA under the project n PRJEB79239.

## AUTHOR CONTRIBUTIONS

CV, JB, PL, TU and GG completed fieldwork and collected samples from Fairbanks, Alaska, USA. MW and SZ screened soil samples for bacterial isolation. MW, SZ performed molecular analysis and MW JB and MV analyzed the sequencing data. BR screened the isolates for EPSs production, MW and JB wrote the manuscript, and CV, GG, HW and OS contributed to and have approved the final manuscript.

