## Supplementary material for "Stick together: Isolation and characterization of exopolysaccharides producing bacteria from degraded permafrost soils": S

### 1. Supplementary figures

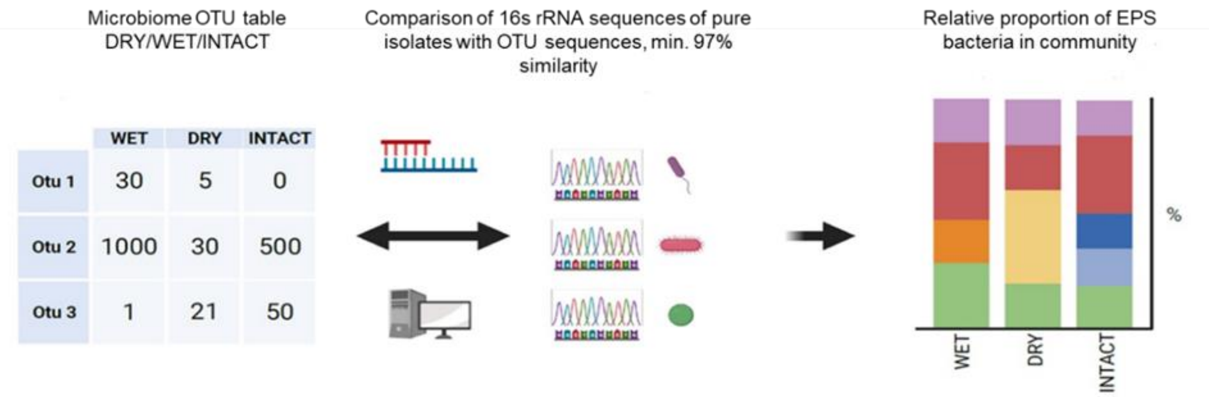


**Figure S1.** Pipeline for the determination of relative proportion of EPS producing soil isolates in total bacterial community of three studies sites. This figure was created with BioRender.com (2022).

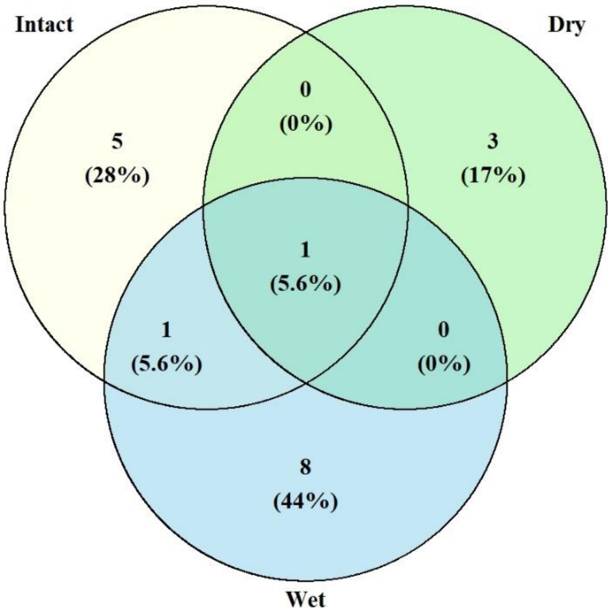


**Figure S2.** This Venn diagram highlighted the diversity and overlap of bacterial isolates across the two sites with degraded permafrost soil (dry and wet sites) and the site with undisturbed permafrost (intact site). Numbers represent the distinct strain count in each section, and percentages are calculated based on the total number of overlapping strains.

**Table S1.** Molecular analysis of soil isolates and their growth on agar medium specific for polysaccharide production. The abbreviations +, ++ and +++ represent the intensity of monoidal/slimy texture of colony. Isolates sharing more than 97% sequence similarity are presented at the species level and lower than 94% are presented at family level.

| **Phylum** | **ID** | **16s rRNA sequencing at species level** | **% Identity** | **Sites** | **PSA Glucose equivalent (mg $L^{-1}$)** | **Mucoid Intensity** |
|---|---|---|---|---|---|---|
| **Firmicutes** | | | | | | |
| | A9 | *Bacillus mycoides* strain ATCC 6462 | 100% | Dry | 68.27 | ++ |
| | A13 | *Bacillus mycoides* strain ATCC 6462 | 100% | Dry | 212.37 | +++ |
| | S28 | *Bacillus mycoides* strain ATCC 6462 | 100% | Dry | 140.79 | ++ |
| | A21 | *Peribacillus simplex* strain LMG 11160 | 99.9% | Dry | 82.87 | + |
| | S27 | *Peribacillus simplex* strain LMG 11160 | 99.6% | Dry | 96.95 | ++ |
| | A3 | *Viridibacillus arvi strain* LMG 22165 | 100% | Dry | 43.53 | ++ |
| | A4 | *Viridibacillus arvi strain* LMG 22165 | 99.4% | Dry | *NA* | +++ |
| | S11 | *Viridibacillus arvi strain* LMG 22165 | 99.4% | Dry | 6.41 | + |
| | S24 | *Viridibacillus arvi strain* LMG 22165 | 100% | Dry | 8.75 | ++ |
| | S30 | *Viridibacillus arenosi* strain LMG 22166 | 99.4% | Dry | 5.38 | + |
| | S34 | *Paenibacillus borealis* strain KK19 | 98.3% | Dry | 46.14 | + |
| | S13 | *Staphylococcus epidermidis* strain Fussel | 99.8% | Dry | *NA* | + |
| | A19 | *Bacillus subtilis* strain JCM 1465 | 99.8% | Intact | 92.9 | +++ |
| | S29 | *Bacillus mycoides* strain ATCC 6462 | 99.9% | Intact | 145.34 | +++ |
| | S31 | *Bacillus mycoides* strain ATCC 6462 | 100% | Intact | 180.69 | +++ |
| | S37 | *Bacillus mycoides* strain ATCC 6462 | 97.6% | Intact | 150.05 | ++ |

| | | | | | | |
|---|---|---|---|---|---|---|
| | A6 | *Bacillus proteolyticus* strain MCCC 1A00365 | 98.0 | Intact | 170.17 | ++ |
| | S35 | *Domibacillus mangrovi* strain SAOS 44 | 98.7% | Intact | 43.54 | + |
| | A20 | *Tumebacillus permanentifrigoris* strain Eur1 9.5 | 99.4% | Intact | *NA* | ++ |
| | A7 | *Staphylococcus saprophyticus* subsp. *saprophyticus* ATCC 15305 = NCTC 7292 | 94.1% | Intact | *NA* | +++ |
| | A24 | *Bacillus mycoides* strain ATCC 6462 | 100% | Wet | 515.24 | ++ |
| | S33 | *Bacillus mycoides* strain ATCC 6462 | 99.9% | Wet | 54.51 | + |
| | S38 | *Bacillus subtilis* strain SBMP4 | 99.1% | Wet | 171.48 | ++ |
| | S39 | *Bacillus subtilis* strain SBMP4 | 98.0% | Wet | 164.54 | ++ |
| | A25 | *Neobacillus bataviensis* strain NBRC 102449 | 99% | Wet | 1148.6 | +++ |
| | S19 | *Neobacillus pocheonensis strain Gsoil 420* | 99.1% | Wet | 146 | ++ |
| | S3 | *Peribacillus simplex* strain LMG 11160 | 96.2% | Wet | 100 | ++ |
| | S4 | *Peribacillus simplex* NBRC 15720 = DSM 1321 | 95.5% | Wet | 86.61 | ++ |
| | S6 | *Mesobacillus subterraneus* strain COOI3B | 98.2% | Wet | 911.44 | *X* |
| | A8 | *Staphylococcus saprophyticus* subsp. *saprophyticus* ATCC 15305 = NCTC 7292 | 100% | Wet | *NA* | + |
| **Actinomycetota** | | | | | | |
| | A11 | *Streptomyces brevispora* strain BK160 | 99.6% | Dry | 342.92 | +++ |
| | S14 | *Streptomyces luozhongensis* strain TRM49605 | 99.5% | Dry | *NA* | + |
| | S21 | *Streptomyces luozhongensis* strain TRM49605 | 99.2% | Dry | *NA* | + |
| | A2 | *Microbacterium flavescens* strain 401 | 98.3% | Dry | 136.85 | + |
| | S17 | *Frigoribacterium faeni* strain 801 | 99.4% | Intact | 942.57 | ++ |
| | S18 | *Frigoribacterium faeni* strain 801 | 99.5% | Intact | 735.79 | +++ |
| | A17 | *Curtobacterium oceanosedimentum* strain ATCC 31317 | 99.3% | Intact | 1309.14 | +++ |
| | S15 | *Pseudarthrobacter sulfonivorans* strain ALL | 98.6% | Intact | 115.63 | +++ |
| | S23 | Micrococcaceae | 91.7% | Intact | 208.21 | ++ |
| | S9 | *Streptomyces luozhongensis* strain TRM49605 | 99.5% | Wet | 118.63 | + |
| | S12 | *Streptomyces laculatispora* strain BK166 | 99.5% | Wet | 3.16 | + |
| | A18 | *Micromonospora fulva* strain UDF-1 | 98.6% | Intact | *NA* | + |
| | S7 | *Curtobacterium oceanosedimentum strain ATCC 31317* | 98.8% | Wet | 216.69 | +++ |
| | S10 | *Micrococcus yunnanensis* strain YIM 65004 | 99.3% | Wet | 22.27 | + |
| **Pseudomonadota** | | | | | | |
| | A12 | *Luteimonas arsenica strain* 26-35 | 99.8% | Dry | 227.26 | ++ |
| | A22 | *Phyllobacterium loti strain S658* | 99.0% | Wet | 177.28 | ++ |
| | A23 | *Phyllobacterium endophyticum* strain PEPV15 | 99.5% | Wet | 124.15 | ++ |
| | S5 | Enterobacteriaceae | 91.2% | Wet | 667.23 | + |
| **Unidentified** | | | | | | |
| | A15 | Unsequenced | | Intact | 126.74 | + |
| | A16 | Unsequenced | | Intact | 35.72 | + |

| | | | | |
|---|---|---|---|---|
| S1 | Unsequenced | Wet | 56.39 | + |
| S2 | Unsequenced | Wet | 966.25 | +++ |
| S8 | Unsequenced | Wet | 124.75 | ++ |
| S20 | Unsequenced | Wet | 157 | + |

**Table S2.** EPSs-producing isolates sequencing similarities to 16S rRNA sequences of zOTUs from the total bacterial community. The threshold from similarity between sequences was set at greater than 97%.

| **Phylum** | **Soil isolates** | **Assigned zOTUs** | **No. of soil isolates with assigned zOTUs** | **% Identity** |
|---|---|---|---|---|
| **Firmicutes** | | | | |
| | *Mesobacillus subterraneus* | *Mesobacillus*-zOTU3623 | 1 | 98.1% |
| | *Neobacillus bataviensis* | *Neobacillus*-zOTU702 | 1 | 100% |
| | *Neobacillus pocheonensis* | *Neobacillus*-zOTU702 | 1 | 100% |
| | *Bacillus mycoides* | *Bacillus*-zOTU2010 | 6 | 98.1% |
| | *Bacillus subtilis* | *Bacillus*-zOTU3 | 2 | 100% |
| | *Peribacillus simplex* | *Peribacilus*-zOTU2176 | 1 | 99.8% |
| **Actinomycetota** | | | | |
| | *Frigoribacterium faeni* | *Frigoribacterium*-zOTU1136 | 2 | 99.6% |
| | *Curtobacterium oceanosedimentum* | *Curtobacterium*-zOTU2238 | 2 | >98.8% |
| | *Streptomyces luozhongensis* | *Streptomyces*-zOTU1852 | 1 | 98.8% |
| | *Streptomyces brevispora* | *Streptomyces*-zOTU1848 | 1 | 97.7% |
| | *Pseudarthrobacter sp.* | *Pseudarthrobacter*-zOTU1072 | 2 | 100% |
| | *Microbacterium flavescens* | *Microbacterium*-zOTU1055 | 1 | 97.7% |
| **Pseudomonadota** | | | | |
| | *Luteimonas arsenica* | *Luteimonas*-zOTU2611 | 1 | 98.5% |
| | *Phyllobacterium loti* | *Phyllobacterium*-zOTU3875 | 1 | 100% |
| | *Phyllobacterium endophyticum* | *Phyllobacterium*-zOTU3875 | 1 | 100% |
